# Gcn4 regulates phospholipid methylation and SAM dependent methyl allocations between phospholipids and histones during methionine-supplemented growth

**DOI:** 10.1101/2025.07.01.662382

**Authors:** Swati Shyam Prasad, Gowda Sampreeth Ravi, Vandana Sah, Vineeth Vengayil, Sunil Laxman, Rajalakshmi Srinivasan

**Affiliations:** BRIC Institute for Stem Cell Science and Regenerative Medicine, Bengaluru – 560065, India; Institute of Bioinformatics and Applied Biotechnology, Bengaluru – 560100, India; Manipal Academy of Higher Education, Manipal – 576104, India

## Abstract

Methionine availability alters the methylation status of various cellular molecules, and thereby controls metabolic state, via its metabolite S-adenosyl methionine (SAM). In yeast cells, supplementing excess methionine drives cell growth and proliferation, as well as phospholipid biogenesis. High methionine stabilizes the transcription factor Gcn4 (Atf4), which drives an anabolic transcription program to sustain high growth. Here, we investigated whether Gcn4 regulates the allocation of SAM towards distinct methylation outputs, during this methionine-dependent anabolic program. We find that methionine induced Gcn4 increases the expression of phospholipid methyltransferases during methionine supplementation. In *Δgcn4* cells, the expression of these methyltransferases significantly decreases, and results in reduced SAM allocations towards phospholipid methylation, alongside increased intracellular SAM, disrupting the balance in the SAM/SAH ratio in these cells. Excess SAM pools are re-allocated to histones, resulting in their hyper-methylation. Collectively, we uncover that Gcn4 modulates SAM allocations towards phospholipid methyltransferases, and the overall methyl allocation between phospholipids and histones. Our study reveals a novel role for Gcn4 in SAM dependent phospholipid biosynthesis in cells growing in methionine sufficiency.

## Introduction

Methionine is not merely another amino acid and signaling molecule, but is the primary precursor of the main biological methyl donor, S-adenosyl methionine (SAM) [1–6]. SAM donates its methyl group to various biological molecules including proteins, DNA, RNA and phospholipids [5,7]. Methylation of these molecules is facilitated by the conversion of SAM to S-adenosyl homocysteine (SAH) [5,7]. As SAH accumulates, product-based feedback inhibits methyltransferases, and cells thereby maintain their internal SAM/SAH ratio [4,5,8,9]. Studies in model cells including yeast have revealed a mechanistic understanding of the role of methionine and SAM in driving growth programs, as well as identified mechanisms through which the methionine dependent growth is mediated [3,10–12]. When methionine, and thereby SAM levels increase, cell switch on strong growth programs, which are built concurrently around high reductive biosynthetic capacity, increased amino acid production, ribosomal biogenesis, the activation of TORC1, and increased phospholipid synthesis [1,2,6,9–14]. Under methionine supplemented growth, the increased methylation of the protein phosphatase PP2A regulates the phosphorylation status of the downstream effectors such as the Gcn4 transcription factor, or the TORC1 regulator Npr2, which subsequently regulates growth and autophagy, respectively [3,11]. On the other hand, under methionine depleted conditions, the reduced methylation of PP2A leads to demethylation of histones, mediated by the phosphorylation status of the histone demethylases [8]. Prior studies now reveal a close coupling between altered histone methylation and phospholipid methylation [8,9,15–17]. Overall, altered SAM levels, under methionine supplemented or the depleted conditions, mediates opposite cellular processes primarily by altering the relative methylation status of proteins, histones and phospholipids, which in turn reflects on the gene regulation, metabolism and eventually on the cell fate [2,3,9,11,16,18,19].

In fast growing cells, a significant proportion of SAM is consumed in phospholipid methylation – a key reaction in phospholipid biosynthesis [9,13]. Trimethylation of the phospholipid phosphatidylethanolamine (PE) leads to the synthesis of phosphatidylcholine (PC). PC is the most abundant phospholipid in the cell [20–23]. In the absence of free choline, phosphatidylethanolamine methylation is the primary pathway towards *de novo* PC biosynthesis, and a large proportion of the methyl pools is consumed during PE methylation [9,15,24]. These methylation reactions are catalysed by phosphatidylethanolamine methyltransferases (PEMTs) such as Cho2 and Opi3 [9,15,25–27]. Cho2 is a methyltransferase that catalyses the first step of phosphatidylethanolamine trimethylation, and the downstream methylation reactions are catalysed by Opi3 [27]. Expression and the activities of these phospholipid methyltransferases are responsive to the SAM levels [9,13,15,24]. Cells lacking PE methyltransferases Cho2 accumulate SAM, indicating the role of Cho2 in balancing the internal SAM/SAH ratio [9,15,19]. Overall, inhibiting phospholipid methylation leads to the accumulation of SAM and the reduced production of SAH, which in turn leads to a skewed SAM/SAH ratio [9,13,24]. In cells defective for phospholipid methylation, excess SAM is diverted to histones, leading to their hypermethylation, thereby altering the epigenetic status of the cell [9,19]. The sequential flow of methyl groups to different methylation sites on histone has been reported [9]. These studies suggest that a controlled SAM allocation to different molecules is mandatory in order to maintain the SAM/SAH ratio and cellular homeostasis. Overall, the methylation status of different molecules in the cell, optimal SAM/SAH ratios, and the regulation of methyltransferases are closely coupled [3,8,9,11,17]. Therefore, asking what controls the allocation of SAM towards the methylation of distinct molecules under conditions of SAM excess (such as during methionine supplemented conditions) becomes important question to be addressed.

In yeast cells growing in high methionine, we uncovered that the amounts of the Gcn4 transcription factor substantially increases [2,12]. Gcn4 is a master regulator of amino acid biosynthesis best studied under starvation, but is also required to sustain amino acid synthesis during high growth [2,11,12,28–30]. Gcn4 levels increase in cells supplemented with methionine, via the stabilization of this protein due to the specific activity of methylated PP2A, and is crucial for sustaining the anabolic program during methionine-supplemented growth [1,2,12]. *Δgcn4* cells in excess methionine show slow growth, and a collapse of the anabolic transcriptional program, reiterating the importance of Gcn4 in driving growth during methionine supplemented conditions [2,12].

In this study, we asked if Gcn4 has any role in modulating the substantial allocations of methylation of distinct molecules, and thereby linking SAM utilization with the overall anabolic response in methionine supplemented condition. We uncover that in methionine-supplemented conditions, Gcn4 play a crucial role in balancing the SAM allocations between phospholipid synthesis and histones, primarily by regulating the genes involved in phospholipid methylation. The loss of Gcn4 results in a sharp downregulation of genes involved in phospholipid methylation, leading to the accumulation of excess SAM. Further, the excess SAM pools are diverted towards increased histone methylation, resulting in histone hypermethylation in *Δgcn4*. We therefore uncover a novel role of Gcn4 in balancing SAM allocations between phospholipids and histones, primarily by regulating phospholipid methyltransferase expression under methionine-supplemented growth conditions.

## Results

### Gcn4 drives the expression of phospholipid methyltransferases during methionine-supplemented growth

Phosphatidylcholine (PC) is the most abundant phospholipids in fast growing cells, and is primarily synthesised as a product of the trimethylation of phosphatidyl ethanolamine (PE) [13,21,31], consuming three methyl groups for the synthesis of each PC molecule. In *Saccharomyces cerevisiae*, this methylation of PE is catalysed by the PE methyltransferases Cho2 and Opi3 [9,15,21,32]. In cells grown in excess of methionine, the Cho2 levels increase, to sustain the high phospholipid biosynthesis required [1,2,9,12,13,19,32]. Separately, Gcn4 was found to be crucial for driving the overall transcriptional program to sustain high amino acid biosynthesis, in these conditions [2,11,12]. We therefore asked if Gcn4 was also involved in this process of PE methylation and PC biosynthesis. We carefully analyzed transcriptomes of wild-type and *Δgcn4* cells grown in high methionine [12]. Notably, the transcripts of enzymes required for phosphatidylcholine synthesis, specifically Opi3 and Cho2 that code for PE methyltransferases that synthesise PC were substantially downregulated in *Δgcn4* (Figure 1a). This revealed that Gcn4 was necessary for sustaining the enzyme levels required for phospholipid methylation, during methionine-supplemented growth (Figure 1a).

**Figure 1:**
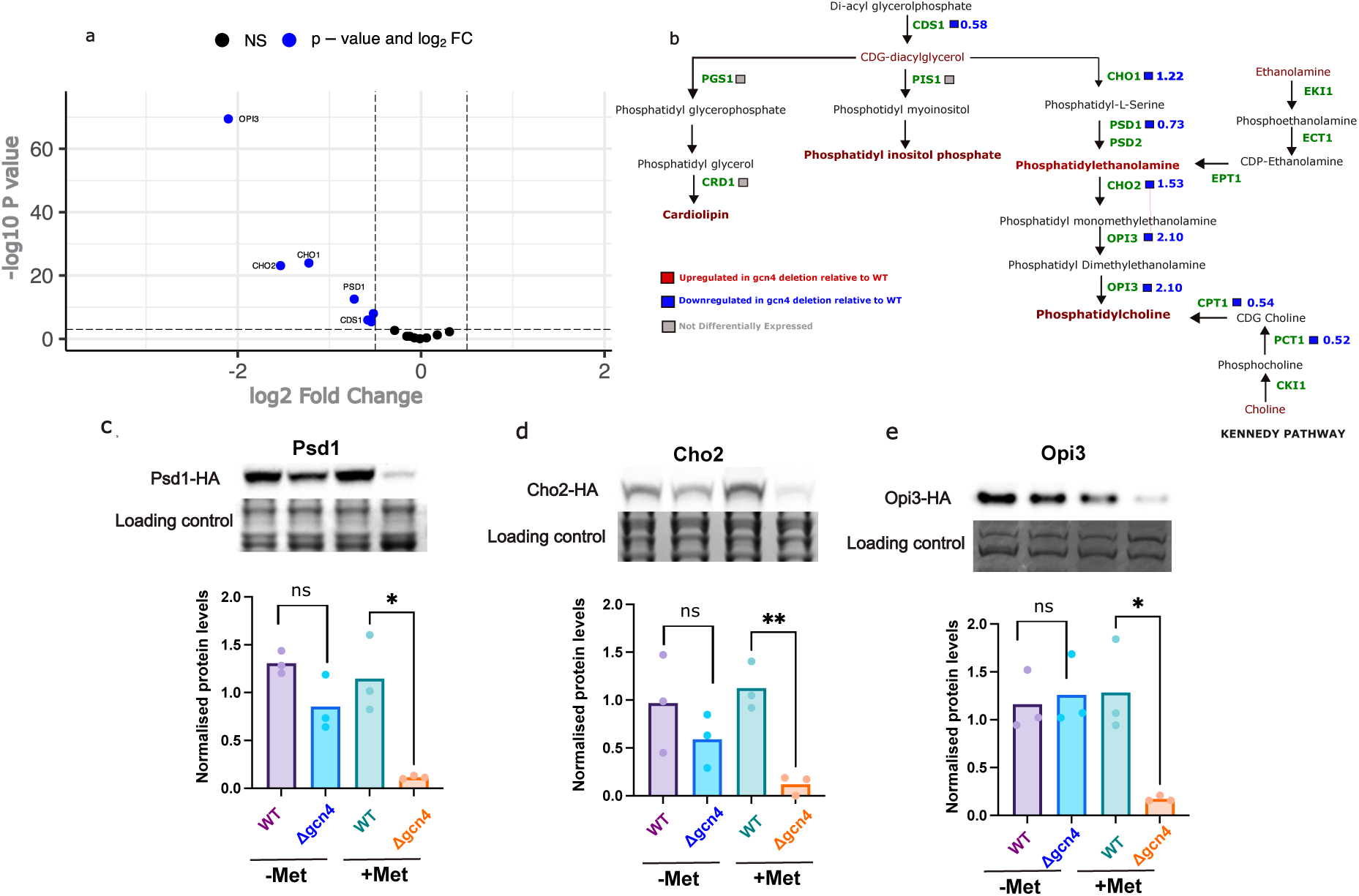
Gcn4 positively regulates genes required for phospholipid methylation in methionine supplemented media. a. Volcano plots showing the differential expression of the phospholipid biogenesis genes in Gcn4 deletion (Δgcn4) relative to the wild-type cells in +Met condition. The x-axis shows the Log2-fold change and the Y-axis shows −log10(P-values) for the genes involved in yeast’s phospholipid biogenesis pathways. Refer to Supplementary Sheet S1. b. Log2 FC values of the genes involved in the given pathway are mapped on the superpathway of the phospholipid biosynthesis pathway map. Blue color boxes represent the given gene is downregulated in Δgcn4 relative to WT. Red and Grey boxes show upregulation and non-differentially expressed genes, respectively. None of the genes belonging to this superpathway are upregulated in Δgcn4. c. Western blot quantification of the protein Psd1 – a protein involved in phospholipid biogenesis. d,e. Western blotting quantification of the phospholipid methyl transferases Cho2 and Opi3, respectively. Bar plots show normalised protein abundance of the given protein with individual data points for 3 replicates. Student t-test was used for the statistical analysis * < 0.05, ** < 0.01 and ns – not significant. Coomassie-stained PAGE cells were used as a protein-loading control (total protein). +Met and −Met denote the presence and the absence of methionine supplementation.

We therefore more extensively assessed if Gcn4 broadly controlled the transcription of genes involved in phospholipid biogenesis. To address this, we listed all genes involved in the superpathway of phospholipid biogenesis from yeast pathways [33]. Out of 15 genes belonging to the superpathway of phospholipid biogenesis, 7 were significantly downregulated in the cells lacking Gcn4 (*11gcn4*) (with a minimum of 45% decrease in transcript level relative to wild type) (Figure 1b). Out of these 7 genes, 4 of them (PSD1, CHO1, CHO2, OPI3) are exclusively involved in the PC biosynthetic arm of the superpathway, and are downregulated by ∼1.7 to 4-fold in *Δgcn4* relative to WT cells (Figure 1b), two out of 3 genes CPT1 and PCT1, involved in Kennedy pathway were down regulated by ∼1.4 fold in *11gcn4* cells relative to the WT cells (Figure 1b). Together, these results suggest that the genes involved in phosphatidylcholine biogenesis are severely downregulated in *11gcn4* cells in methionine-supplemented medium.

Given this observed decrease in PC related transcripts, we assessed the respective protein levels of these genes in media with (+Met) or without (-Met) supplementing methionine. Western blot analysis revealed that the levels of Opi3, Cho2 and Psd1 proteins involved in PC biosynthetic pathways decrease substantially, and this was especially apparent for the PE methyltransferases (Opi3 and Cho2) in *Δgcn4* in +Met (Figure 1c,d & e). These data reveal that Gcn4 is required for the expression of the genes involved in phospholipid methylation process in methionine-supplemented conditions (+Met). Note that this effect was pronounced only when the *11gcn4* cells were grown in the methionine-supplemented (+Met) medium, suggesting that this requirement for Gcn4 was methionine-specific.

### Gcn4 regulates phospholipid methyltransferase expression in +Met to a similar extent as Ino2

Ino2 is a well-known, general transcriptional regulator of the PE methyltransferases [34–36]. Earlier studies have established a housekeeping role of the Ino2 transcription regulator in the maintenance of phospholipid biosynthesis, where Opi3 and Cho2 expression is increased due to the Ino2-Ino4 complex binding to their promoters, and the levels of Opi3 and Cho2 transcripts decrease in *Δino2* cells [34,35]. Our data unexpectedly revealed a necessary role for Gcn4 in the expression of the phospholipid methyltransferases Cho2 and Opi3 during methionine supplementation. We therefore asked if Gcn4 mimics the regulatory effect of Ino2, in regulating phospholipid methyltransferases. To address this question, we compared the protein levels of Opi3 and Cho2 in the *Δgcn4, Δino2 and Δgcn4/Δino2* double mutants, grown in the presence of methionine. Interestingly, we observed > 70% reduction of Opi3 and > 98% reduction of Cho2 protein levels in *Δgcn4* cells compared to WT. We also observed > 98% reduction of both Opi3 and Cho2 in the *Δino2 and Δgcn4/Δino2* double mutants (Figure 2a &b), These data suggest that the loss of Gcn4 (*Δgcn4)* alone reduces Cho2 and Opi3 expression to an extent comparable to that of the loss of Ino2 (*Δino2*). This observation reveals an unforeseen role of Gcn4 in regulating phospholipid methylation during methionine supplementation. The role of Gcn4 in controlling the expression of Cho2 and Opi3 is comparable to the effect of the known, housekeeping transcriptional regulator of PL biosynthesis, Ino2 (Figure 2a and b).

**Figure 2:**
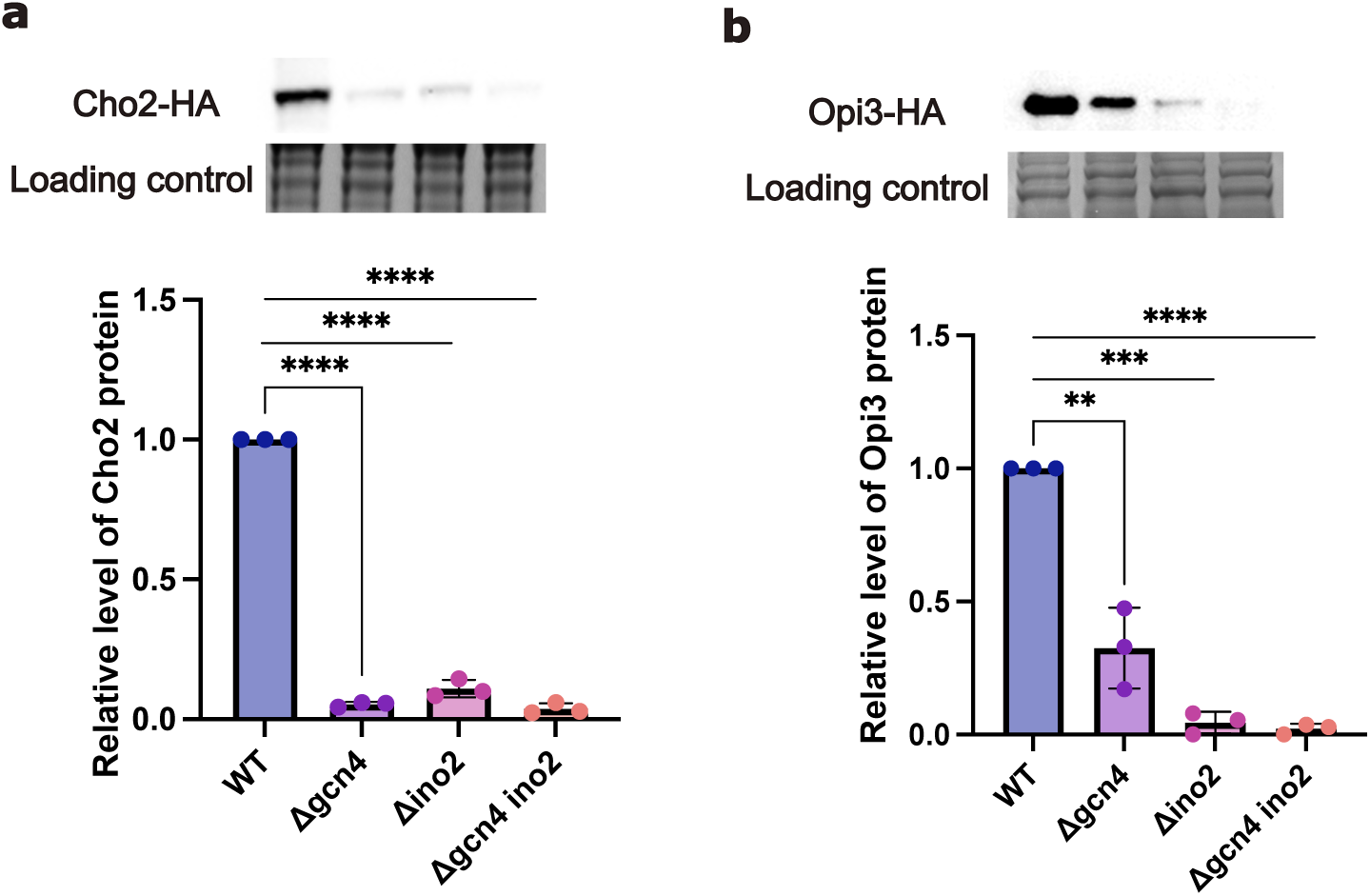
Loss of Gcn4 regulates phospholipid methyltransferases during methionine supplementation to levels comparable with the loss of Ino2. a-b. Western blot and quantification of the Cho2 and Opi3 proteins, respectively, in the given deletion backgrounds. The Cho2 and Opi3 genes were carboxy-terminally tagged at their endogenous locus with a HA-epitope in WT, Δgcn4, Δino2 and Δgcn4/Δino2 strains. Relative protein level of Cho2 and Opi3 normalised to the WT cells are plotted. The cells were grown in the presence of 2mM methionine. Figure shows the reduced level of Cho2 and Opi3 protein in all the above deletion strains of Gcn4 and Ino2, indicating an essential role of Gcn4 in phospholipid biogenesis. Barplots with individual data points are shown. Student t-test was used for the statistical analysis *< 0.05, ***< 0.001, ****< 0.0001 and ns – not significant. Coomassie-stained PAGE cells were used as a protein-loading control.

### Phospholipid levels are decreased in *Δgcn4* cells under methionine supplementation

To investigate whether the loss of Gcn4, and the associated reduction in phospholipid biosynthetic genes, results in changes in total phospholipid amounts, we quantified the levels of major phospholipids using targeted, quantitative mass spectrometry analysis. Compared to the WT cells, *Δgcn4* cells show a substantial reduction (∼75%) in PS (16:1/18:1), ∼44% in PE(16:1/18:1), ∼25% in PC(16:1/18:1), ∼42 % in PS (16:1/16:1) and ∼59% in PE (16:1/16:1) (Figure 3a-c). Collectively, we observed a significant reduction in the levels of phospholipids, particularly in phosphatidylcholine (PC) in *Δgcn4* cells. Given that PC is the most abundant phospholipid, we collectively observe that when methionine levels are high, Gcn4 plays a crucial role in directing the utilization of methyl pools towards PC biogenesis, by sustaining the expression of enzymes leading to the final formation of PC. Interestingly, in media conditions without the methionine supplementation (-Met), there was no significant difference in the phospholipid levels between WT and *Δgcn4* mutants. These data collectively establish a methionine-specific role for Gcn4 in sustaining phospholipid biosynthesis under high methionine conditions.

**Figure 3:**
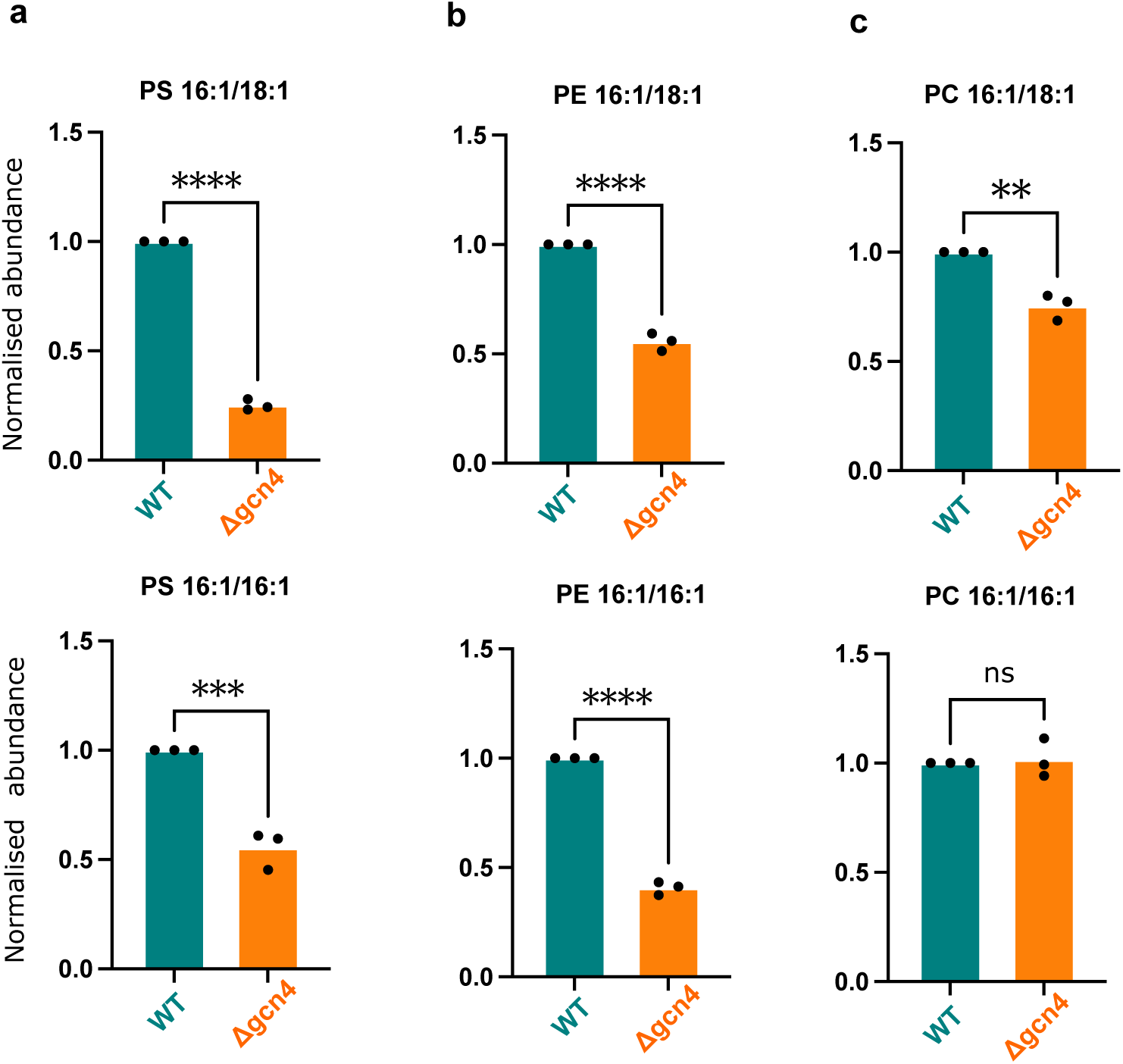
Phospholipid levels are decreased in Δgcn4 cells under methionine supplementation : Steady-state levels of different phospholipids are measured using mass spectrometry. PC and PE are the most abundant phospholipids in yeast cells, accounting for over 80% of total phospholipids. Based on the two most common species of PC and PE, the precursor PS was chosen to represent the reduced levels of phospholipids in the phospholipid biosynthetic pathway. The phospholipids are represented from left to right in order of reactions in the pathway. a. Relative levels of PS 16:1/18:1 and 16:1/16:1. b. Relative levels of PE 16:1/18:1 and PE 16:1/16:1. c. Relative levels of PC 16:1/18:1 and PC 16:1/16:1; a, b, c show a decrease in major phospholipids in the cell and hampered biosynthesis. Unpaired Student t-test was used for the statistical analysis * < 0.05, ** < 0.01, *** < 0.001, **** < 0.0001 and ns – not significant. Data was normalized by accounting for extraction efficiency across samples using an internal. All values were normalised to the WT cells grown in +Met condition and plotted the relative abundance of phospholipids.

### SAM/SAH ratio increases in *11 gcn4* alongside reduced PC levels

In growing cells, phospholipids are a major consumer of SAM, and the loss of phospholipid methyltransferases leads to accumulation of SAM [9,15]. Our data shows reduced synthesis of phospholipid methyltransferases in the *Δgcn4* during methionine-supplemented growth, hence we hypothesised that SAM levels in these cells might increase consequentially (Figure 4a). To investigate this, we measured the steady-state levels of SAM and SAH using mass spectrometry assays in WT and *Δgcn4* cells grown in the presence and absence of methionine. We observed about ∼2-fold increase in SAM amounts in *Δgcn4* mutants compared to WT cells grown in the presence of methionine (+Met) (Figure 4b). The SAH levels between the WT and *Δgcn4* cells were comparable (Figure 4c). These data reveal that SAM accumulates in the *Δgcn4 cells*, leading to an overall increase in SAM:SAH ratio (Figure 4d). This accumulation is specific to the cells grown in the presence of methionine (Figure 4). In the absence of supplemented methionine, there was no significant difference in SAM, SAH and SAM:SAH ratio observed between WT and the *Δgcn4* cells (Figure 4d), reiterating a Gcn4-dependent role in driving SAM consumption in methionine-supplemented cells. This result reveals an imbalance in SAM utilisation in *Δgcn4 cells*, during methionine-supplemented growth. This is also consistent with the substantial reduction in phospholipid amounts in *11gcn4* cells under these conditions.

**Figure 4:**
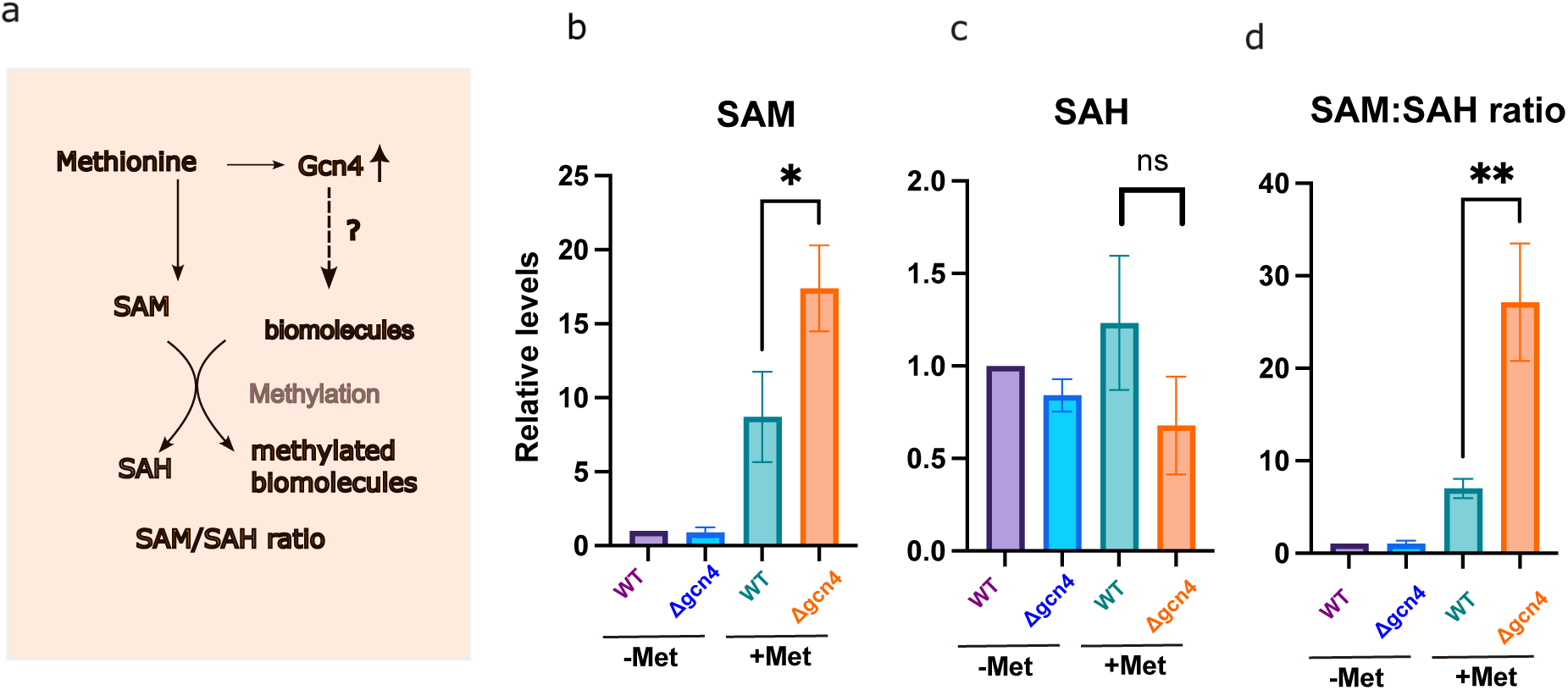
SAM amounts and the SAM/SAH ratio increase in the Δgcn4 cells during methionine supplemented growth: a. Schematic illustrating the hypothesis of the role of Gcn4 in regulating the methylation status of distinct biomolecules (phospholipids, histone proteins), when Gcn4 protein levels increase during methionine supplementation. b. The bar graph show steady state levels of S-Adenosyl Methionine in Saccharomyces cerevisiae cells grown in minimal media in the presence (+Met) or absence of methionine (-Met). WT and Δgcn4 represent the WT and the deletion mutants of Gcn4. The data represented are from minimum of 3 biological replicates. The mean and the SD are shown in the plot. (**p value < 0.005, * p-value < 0.05, ns – non significant – Unpaired Student’s t-test). c. same as 1b, for SAH. d. same as 1b, for SAM:SAH ratio.

### Excess SAM pools are diverted towards histone methylation in *Δgcn4* cells

Prior studies have established that lipid synthesis is a major consumer of SAM (during methionine supplemented conditions) [9], and concurrently, histones serve as major sinks for methylation under these conditions [9,15,16,19]. Since phospholipid methylation is substantially reduced in the *Δgcn4* cells, and the SAM/SAH ratio is high in these cells, we asked if the resultant, excess SAM was diverted towards the methylation of histones (Figure 5a). Towards addressing this, we measured total histone methylation by western blotting using antibodies specific to the major H3K4me3 and H3K36me3 methylation marks. Notably, we observed a substantial increase in the methylated histone levels in *Δgcn4 cells* (Figure 5b and c), while total H3 levels did not change (Figure 5d). These data established that histones were hypermethylated in the *Δgcn4* cells, upon methionine supplementation.

**Figure 5.**
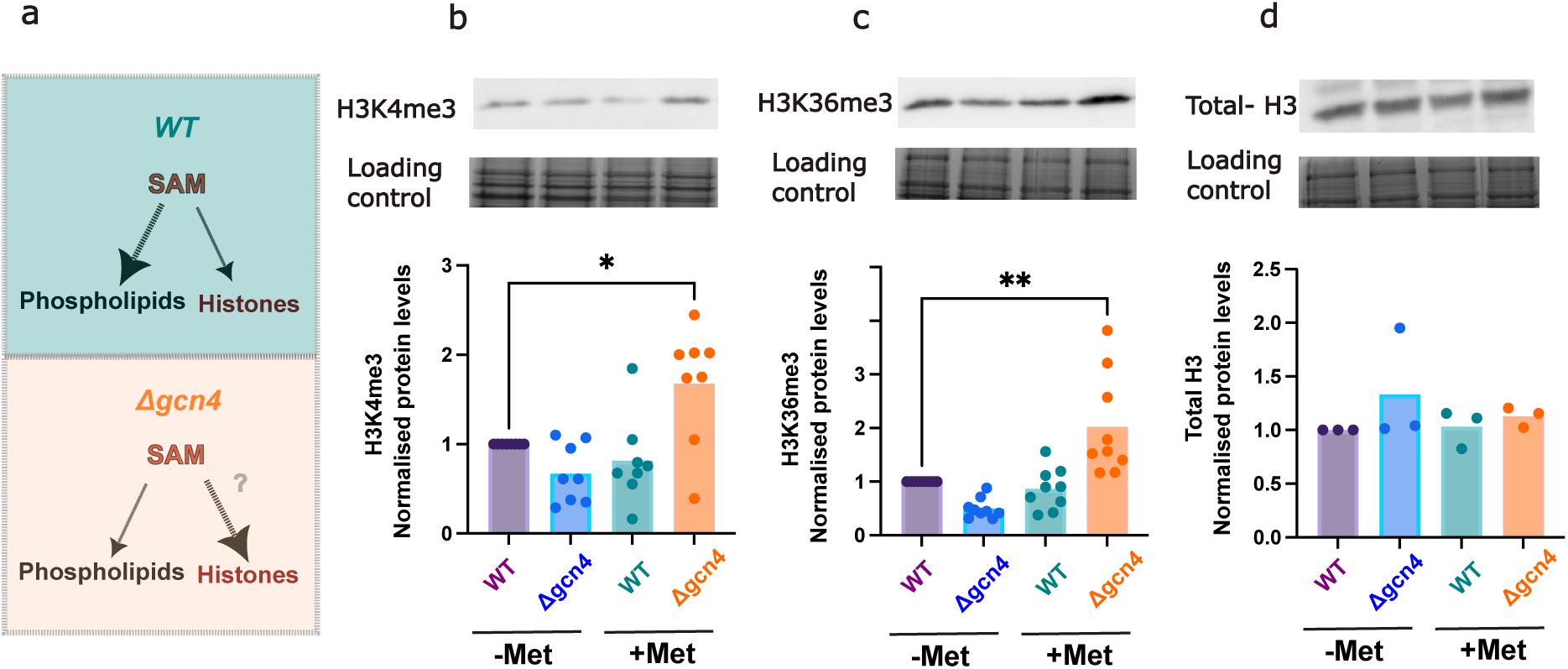
Histone methylation increases in Δgcn4 cells during methionine supplementation: The schematic shows our hypothesis of Gcn4 playing a role in allocating the SAM pools between phospholipids and histone. b-d. Western blot analysis showing the quantification of methylated histones using antibodies specific to H3K4me3, H3K36me3 marks and total H3 proteins. Total protein stained with Coomassie was used as a loading control. Quantifications were relative to wild-type cells grown in the respective media conditions.

Further, since these histones are hypermethylated in *Δgcn4* cells, we also asked if their histone methyltransferases were consequently differentially regulated in these cells. Set1 and Set2 are major histone methyltransferases drive histone methylation at H3K4me3 and H3K36me3 respectively, and other proteins collectively called as COMPASS complexes play a role in this process [37–41]. Existing transcriptome data from cells grown with supplemented methionine, and lacking Gcn4, show a significant upregulation of the histone methyltransferase Set2, as well as several proteins of the COMPASS complex (Supplementary Figure 1a and b). These data are entirely consistent with the observed histone hypermethylation in *Δgcn4*.

Overall, our data reveal a central role for Gcn4 in SAM allocations, and balancing SAM/SAH pools in cells supplemented with methionine, by budgeting them between phospholipid and histone methylation. In the absence of Gcn4, this balance is lost, leading to the disruption of phospholipid methylation, which in turn elevates the methyl pools in the cell. This elevated pool of methyl groups can be re-routed to the histones thus changing the epigenetic landscape of the cell.

## Discussion

Gcn4 is a global transcriptional master regulator primarily studied in the context of increased amino acid biosynthetic gene transcription. Previous studies uncovered that Gcn4 levels substantially increase, via protein stabilization, in cells grown in high methionine, and this increase in Gcn4 drives a methionine dependent anabolic transcriptional program [1,2,11,12]. Separately, it is well established that the intracellular SAM pools are high in cells that are supplemented with methionine [3,8,10]. Gcn4 has been shown to regulate several cellular functions beyond its conventional amino acid stress response, and can support high biosynthesis during cell growth (Walvekar et al., 2018; Walvekar & Laxman, 2019). Given the unique context of rapid growth in methionine, driven by high SAM pools, we asked if Gcn4 play any role in the overall allocations and consumption of SAM for lipid biosynthesis and histone methylation, as reported in previous studies [9,16,19]. Our study reveal a novel, essential role for Gcn4 in maintaining the SAM/SAH ratio of the cell, and appropriate SAM allocations towards phospholipid biosynthesis, during the methionine-driven cell growth program, as shown in the model in Figure 6.

**Figure 6:**
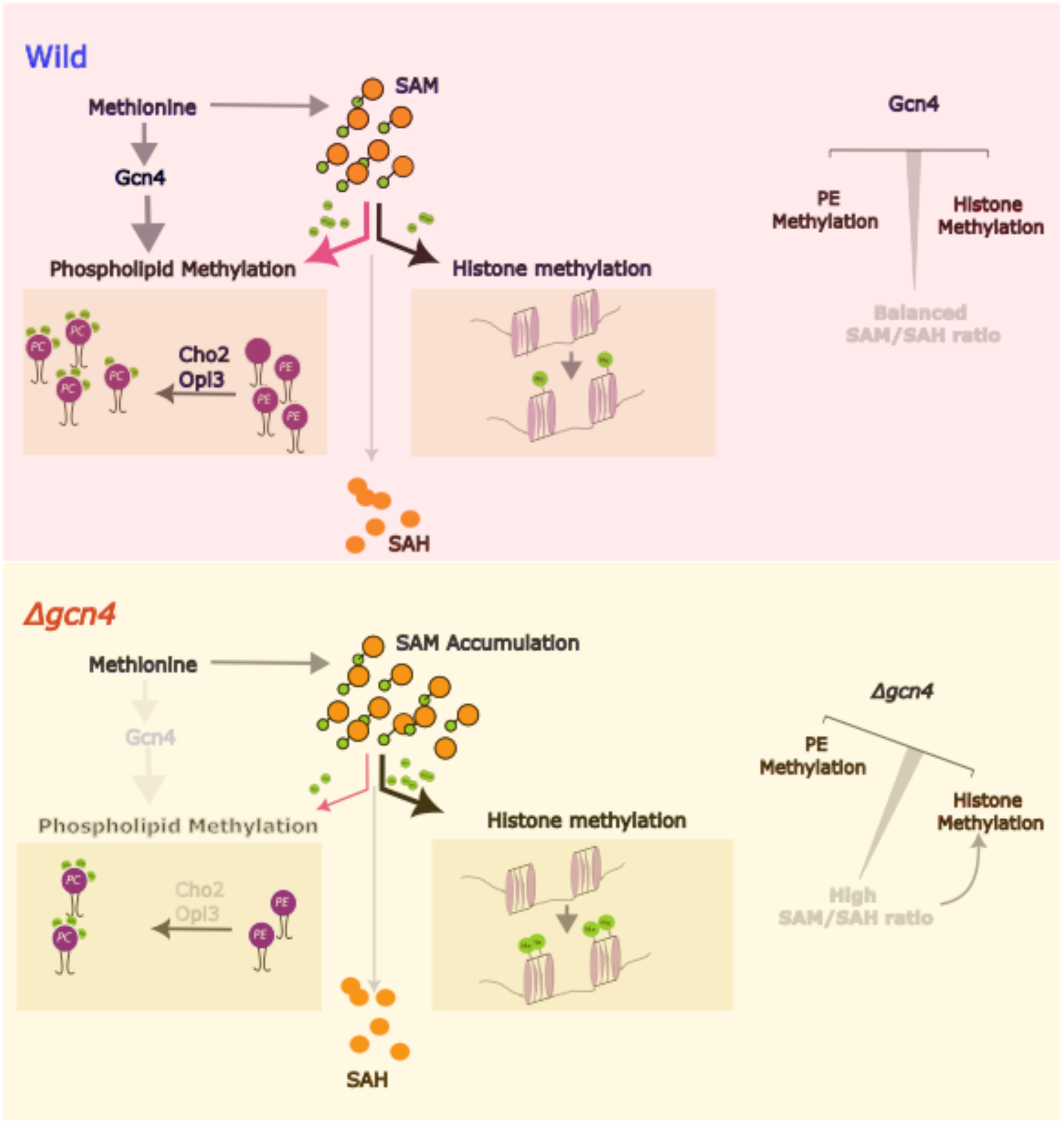
Model depicting the role of Gcn4 in mediating the methyl allocation between phospholipids and histones. This study highlights the role of Gcn4 in balancing SAM/SAH pools by budgeting them to phospholipid and histone methylation. When the WT cells are grown in the presence of methionine, Gcn4 protein level in the cell increases, which activates the Phosphatidylethanolamine methyl transferases(PEMTs), thus keeping the methyl allocation to the phospholipid and histones under control. In the absence of Gcn4, this balance is lost, leading to the disruption of phospholipid methylation, which in turn elevates the methyl pools in the cell. This elevated pool of methyl groups can be re-routed to the histones, thus changing the epigenetic landscape of the cell.

Cells must tightly regulate and maintain their intracellular SAM/SAH ratio (a key indicator of methylation and SAM consumption [4,8,17,42]. How the intracellular SAM/SAH ratio is altered when SAM and methionine are in excess therefore is an important question that we address here. When SAM is abundant, a substantial sink for SAM is phospholipid biosynthesis, given the vast amounts of phospholipids required by cells [9]. Phospholipid methylation is a key step for synthesising the most abundant phospholipid, phosphatidylcholine, [9,13,21,26]. Here, we uncover that when SAM is abundant, Gcn4 drives the methylation of PE, and PC synthesis, by regulating the expression of critical phospholipid methyltransferase genes. Further, under these conditions, Gcn4 drives the upstream biosynthesis of phosphatidyl ethanolamine, and therefore effectively regulates the entire metabolic arm that produces phosphatidylcholine from CDG-2,3-diacylglycerol (Figure 1b). This therefore reveals a central role for Gcn4 in regulating the PC biosynthetic arm, in order to streamline the methylation of phospholipids, during methionine-surplus. Given the absolute requirement for high phosphatidylcholine in cell proliferation and organelle biogenesis [36,43–45], our study reveals how Gcn4 plays a central role in driving a comprehensive growth program in methionine-supplemented conditions, and not merely supporting the required amino acid, nucleotide and ribosomal biogenesis under these conditions.

Furthermore, we uncover that under these conditions, the excess SAM pools that become available in *Δgcn4* cells, are now rerouted to histone methylation. Such a situation will result in a further suppression of growth in *Δgcn4* cells when methionine is abundant [2,12]. These findings reveal a systems-level requirement for cells manage the reciprocal relationship between histone methylation and phosphatidylcholine biosynthesis, where the loss of the PE methylation genes results in histone hypermethylation [17]The role of Gcn4 in mediating this program now becomes comprehensive. Gcn4 regulates amino acid biosynthetic genes, and nucleotide biosynthetic genes, and while some of them are directly regulated by the binding of Gcn4 to its promoter regions, others are indirectly regulated by Gcn4 binding [12]. Some of this might also be a consequence of altered histone methylation observed in *Δgcn4,* explaining the differential gene expression of several indirect targets of Gcn4, for example those reported previously [12]. Overall, our study highlights how a global-transcriptional regulator, Gcn4, balances SAM allocations, by coupling phospholipid metabolism with the histone methylation status of the cell. An obvious line of future enquiry will be to address mechanisms through which cells can synchronise different metabolic outputs with the methylation status of the cell, in order to sustain growth.

## Materials and Methods

### Yeast strains and growth conditions

A prototrophic strain *Saccharomyces cerevisiae CEN.PK* was used for all the experiments [46]. Mutant and chromosomally tagged strains were generated using PCR-based methods. The strain and primer list used in this study are available in the Supplementary Tables 1 and 2. For all the experiments, the overnight-grown primary culture in YPD was shifted to fresh YPD (1% yeast extract, 2% peptone, 2% glucose). Once the OD_600_ of the secondary culture reaches 0.6-0.8, the cells were pelleted down and shifted to the Minimal media (-Met) and Minimal media supplemented with 2mM of methionine (+Met). Cultures were grown in a shaker incubator at 240 rpm, and the temperature was maintained at 30°C.

### Gene Expression Analysis and pathway maps

RNA sequence data from our previous study [12] were used for calling differentially expressed genes between WT and *Δgcn4* mutants grown in the presence of methionine. Differential Gene Expression analysis outputs were listed in Supplementary Sheet S1. The superpathway of phospholipid biosynthesis pathway genes were retrieved from yeast pathways [33]. The list of genes belonging to the superpathway was listed in Supplementary Sheet S1. The Log_2_FC values and adj-p-values for the corresponding genes of the phospholipid biosynthetic pathways were used to plot the volcano plot.

### Western Blot Analysis

The cells grown in YPD were shifted to −Met and +Met conditions. After 1.5 hours of the shift, cells were collected, and total protein was extracted using the Trichloroacetic acid (TCA) method. The protein extracted was resuspended in SDS-Glycerol buffer. Protein samples were run on 4-12% bis-tris SDS-PAGE gels (Invitrogen, NP0336BOX) and transferred to a nylon blotting membrane. A part of the gel was stained with Coomassie blue to normalise for the loading control. Blots were blocked with 5% skimmed milk in 1x TBST, and incubated with primary antibody (1:2000 dilution for anti-HA - Sigma-Aldrich, 11583816001) followed by HRP-labelled secondary antibody,(1:4000 for anti-mouse HRP-labelled antibody, Cell Signalling Technology, 7076S. The blots were developed using ECL-HRP substrate (Advansta, K12045), and chemiluminescence was detected on the chemidoc.

### Western blotting for histone methylation levels

Total protein extraction was done as described in the previous study [47]. Cells corresponding to the OD_600_ of 10, grown in the indicated media were collected by centrifugation at 3500 rpm at 4°C for 5 minutes. The supernatant was removed and the pellet was washed in ice-cold water and centrifuged again. The pellet was resuspended in100 μl of water, 300 μl of 0.2 M sodium hydroxide solution and 20 µl β-mercaptoethanol. The suspension was incubated on ice for 10 minutes and centrifuged at 20,800xg for 10 minutes at 4°C. The pellet was resuspended in 100 µl of SDS-PAGE sample buffer, and boiled at 95°C, for 10 minutes. The samples were then centrifuged at 10,600 × g for 3 min at room temperature, and 6 µl of the supernatant was loaded in a 4-12 % bis tris gel (Invitrogen, cat no: NPO322BOX). The protein was transferred to a PVDF membrane using sodium carbonate transfer buffer (1× sodium carbonate transfer buffer, 20 % methanol), at 22V for 90 minutes at room temperature. The membrane was then blocked in 5% skimmed milk in 1X TBST for 1 hour. The blots were incubated with a 1:2000 dilution of anti-H3K4me3 (abcam, ab12209), anti-H3K36me3 (abcam, ab9050) and anti-histone antibodies (abcam, ab1791) for 2 hours. The membrane was then washed in TBST and incubated with anti-rabbit IgG secondary antibody for 30 minutes (1:4000 dilution). The membrane was washed in TBST and developed using a chemiluminescence reagent (Advansta) and imaged using ChemiDoc. A portion of the gel was stained with Coomassie blue to be used as a total protein control.

### Metabolite Extractions for phospholipid and other metabolite quantifications by targeted LC-MS/MS

The metabolite extraction protocol for SAM, SAH was as done previously [2]. Synergi 4-μm Fusion-RP 80 Å (150 × 4.6 mm) LC column (Phenomenex, 00F-4424-E0) was used to resolve metabolites. Phospholipid extractions and quantifications were adapted from a previous study [48]. Briefly, *S. cerevisiae* cells were grown in the aforementioned media conditions. Yeast cells corresponding to 4 OD were lysed using glass beads in a homogenization solution that constituted of methanol and an internal standard (IS) - to check efficiency of extraction and for quantification. Phospholipids were enriched from this lysate using chloroform. This mixture was vortexed well and centrifuged. To the supernatant, chloroform and citric acid were added, vortexed and centrifuged. Upon centrifugation, phase separation occurred, after which the top and the middle protein interface layer were discarded. The bottom organic phase containing phospholipids was collected and dried using vacuum concentrator. This product was stored at −80°C until further usage. Luna Omega 3 μm Polar C18 100 (100 x 4.6 mm) LC column (Phenomenex, 00D-4760-E0) was used to resolve metabolites. Solvent A - 5mM ammonium acetate in 33.3% methanol, 33.3% acetonitrile and Solvent B - 5mM ammonium acetate in 2-Propanol were used. ABSciex QTRAP 5500 mass spectrometer (with Analyst 1.6.2 software (Sciex)0 was used to detect metabolites. Analysis was done using MultiQuant version 3.0.1 (Sciex). LC-MS/MS parameters for metabolites are in Supplementary Table S3 & S4. Raw data from the mass spec experiments were provided in Supplementary Sheet S2. For normalisation, the area under the peak for the IS is taken for all samples and normalized to a single value. The raw value of the area under the peaks for the rest of the samples are then divided by the normalisation factors, such that extraction discrepancies are corrected for. These corrected values were used for relative quantification.

## Supporting information

Supplementary Sheet S1

Supplementary Sheet S2

## Authors Contribution

RS, SL conceived and conceptualised the study. SP,VV,SG,VS,RS,SL performed the experiments, generated and analysed the data, RS, SL and SP wrote the manuscript.

## Acknowledgement

We acknowledge extensive use of the mass spectrometers at the inStem and NCBS-MS facilities, and the infrastructural facilities at IBAB and inStem.

## Reference

[1] Walvekar AS, Laxman S. Methionine at the Heart of Anabolism and Signaling: Perspectives From Budding Yeast. Front Microbiol 2019. 10.3389/fmicb.2019.02624.

[2] Walvekar AS, Srinivasan R, Gupta R, Laxman S. Methionine coordinates a hierarchically organized anabolic program enabling proliferation. Mol Biol Cell 2018. 10.1091/mbc.e18-08-0515.

[3] Laxman S, Sutter BM, Tu BP. Methionine is a signal of amino acid sufficiency that inhibits autophagy through the methylation of PP2A. Autophagy 2014;10. 10.4161/auto.27485.

[4] Walsh CT, Tu BP, Tang Y. Eight Kinetically Stable but Thermodynamically Activated Molecules that Power Cell Metabolism. Chem Rev 2018;118:1460–94. 10.1021/acs.chemrev.7b00510.

[5] Loenen WAM. S-Adenosylmethionine: jack of all trades and master of everything? Biochem Soc Trans 2006;34:330. 10.1042/BST20060330.

[6] Sutter BM, Wu X, Laxman S, Tu BP. Methionine Inhibits Autophagy and Promotes Growth by Inducing the SAM-Responsive Methylation of PP2A. Cell 2013;154:403–15. 10.1016/j.cell.2013.06.041.

[7] Chiang PK, Gordon RK, Tal J, Zeng GC, Doctor BP, Pardhasaradhi K, et al. S-Adenosylmethionine and methylation. FASEB J 1996;10:471–80.

[8] Ye C, Sutter BM, Wang Y, Kuang Z, Zhao X, Yu Y, et al. Demethylation of the Protein Phosphatase PP2A Promotes Demethylation of Histones to Enable Their Function as a Methyl Group Sink. Mol Cell 2019. 10.1016/j.molcel.2019.01.012.

[9] Ye C, Sutter BM, Wang Y, Kuang Z, Tu BP. A Metabolic Function for Phospholipid and Histone Methylation. Mol Cell 2017. 10.1016/j.molcel.2017.02.026.

[10] Laxman S, Sutter BM, Wu X, Kumar S, Guo X, Trudgian DC, et al. Sulfur Amino Acids Regulate Translational Capacity and Metabolic Homeostasis through Modulation of tRNA Thiolation. Cell 2013;154:416–29. 10.1016/j.cell.2013.06.043.

[11] Walvekar AS, Kadamur G, Sreedharan S, Gupta R, Srinivasan R, Laxman S. Methylated PP2A stabilizes Gcn4 to enable a methionine-induced anabolic program. Journal of Biological Chemistry 2020. 10.1074/jbc.ra120.014248.

[12] Srinivasan R, Walvekar AS, Rashida Z, Seshasayee A, Laxman S. Genome-scale reconstruction of Gcn4/ATF4 networks driving a growth program. PLoS Genet 2020;16:e1009252. 10.1371/journal.pgen.1009252.

[13] Hickman MJ, Petti AA, Ho-Shing O, Silverman SJ, McIsaac RS, Lee TA, et al. Coordinated regulation of sulfur and phospholipid metabolism reflects the importance of methylation in the growth of yeast. Mol Biol Cell 2011;22:4192–204. 10.1091/mbc.e11-05-0467.

[14] Laxman S, Sutter BM, Shi L, Tu BP. Npr2 inhibits TORC1 to prevent inappropriate utilization of glutamine for biosynthesis of nitrogen-containing metabolites. Sci Signal 2014;7. 10.1126/scisignal.2005948.

[15] Strzyz P. Metabolism: Methyl groups sink into phospholipids and histones. Nat Rev Mol Cell Biol 2017;18. 10.1038/nrm.2017.44.

[16] Karimian A, Vogelauer M, Kurdistani SK. Metabolic sinkholes: Histones as methyl repositories. PLoS Biol 2023;21. 10.1371/journal.pbio.3002371.

[17] Fang W, Zhu Y, Yang S, Tong X, Ye C. Reciprocal regulation of phosphatidylcholine synthesis and H3K36 methylation programs metabolic adaptation. Cell Rep 2022;39:110672. 10.1016/j.celrep.2022.110672.

[18] Sutter BM, Wu X, Laxman S, Tu BP. XMethionine inhibits autophagy and promotes growth by inducing the SAM-responsive methylation of PP2A. Cell 2013;154. 10.1016/j.cell.2013.06.041.

[19] Ye C, Tu BP. Sink into the Epigenome: Histones as Repositories That Influence Cellular Metabolism. Trends in Endocrinology & Metabolism 2018;29:626–37. 10.1016/j.tem.2018.06.002.

[20] Acoba MG, Senoo N, Claypool SM. Phospholipid ebb and flow makes mitochondria go. Journal of Cell Biology 2020. 10.1083/JCB.202003131.

[21] Liu YC, Lin YC, Kanehara K, Nakamura Y. A methyltransferase trio essential for phosphatidylcholine biosynthesis and growth 1[OPEN]. Plant Physiol 2019;179. 10.1104/pp.18.01408.

[22] Boumann HA, Gubbens J, Koorengevel MC, Oh CS, Martin CE, Heck AJR, et al. Depletion of phosphatidylcholine in yeast induces shortening and increased saturation of the lipid acyl chains: Evidence for regulation of intrinsic membrane curvature in a eukaryote. Mol Biol Cell 2006;17. 10.1091/mbc.E05-04-0344.

[23] Bürgermeister M, Birner-Grünberger R, Nebauer R, Daum G. Contribution of different pathways to the supply of phosphatidylethanolamine and phosphatidylcholine to mitochondrial membranes of the yeast Saccharomyces cerevisiae. Biochim Biophys Acta Mol Cell Biol Lipids 2004. 10.1016/j.bbalip.2004.09.007.

[24] Sadhu MJ, Moresco JJ, Zimmer AD, Yates JR, Rine J. Multiple inputs control sulfur-containing amino acid synthesis in *Saccharomyces cerevisiae*. Mol Biol Cell 2014;25:1653–65. 10.1091/mbc.e13-12-0755.

[25] Verkerke ARP, Ferrara PJ, Lin C Te, Johnson JM, Ryan TE, Maschek JA, et al. Phospholipid methylation regulates muscle metabolic rate through Ca2+ transport efficiency. Nat Metab 2019;1. 10.1038/s42255-019-0111-2.

[26] Kleetz J, Welter L, Mizza AS, Aktas M, Narberhaus F. Phospholipid N-Methyltransferases Produce Various Methylated Phosphatidylethanolamine Derivatives In Thermophilic Bacteria. Appl Environ Microbiol 2021;87. 10.1128/AEM.01105-21.

[27] Kodaki T, Yamashita S. Yeast phosphatidylethanolamine methylation pathway. Cloning and characterization of two distinct methyltransferase genes. J Biol Chem 1987;262:15428–35.

[28] Natarajan K, Meyer MR, Jackson BM, Slade D, Roberts C, Hinnebusch AG, et al. Transcriptional Profiling Shows that Gcn4p Is a Master Regulator of Gene Expression during Amino Acid Starvation in Yeast. Mol Cell Biol 2001. 10.1128/mcb.21.13.4347-4368.2001.

[29] Hinnebusch AG. Translational regulation of yeast GCN4: A window on factors that control initiator-tRNA binding to the ribosome. Journal of Biological Chemistry 1997. 10.1074/jbc.272.35.21661.

[30] Gupta R, Adhikary S, Dalpatraj N, Laxman S. An economic demand-based framework for prioritization strategies in response to transient amino acid limitations. Nat Commun 2024;15:7254. 10.1038/s41467-024-51769-w.

[31] Li J, Xin Y, Li J, Chen H, Li H. Phosphatidylethanolamine N-methyltransferase: From Functions to Diseases. Aging Dis 2023;14. 10.14336/AD.2022.1025.

[32] Summers EF, Letts VA, McGraw P, Henry SA. Saccharomyces cerevisiae cho2 mutants are deficient in phospholipid methylation and cross-pathway regulation of inositol synthesis. Genetics 1988;120. 10.1093/genetics/120.4.909.

[33] Cherry JM, Adler C, Ball C, Chervitz SA, Dwight SS, Hester ET, et al. SGD: Saccharomyces genome database. Nucleic Acids Res 1998. 10.1093/nar/26.1.73.

[34] Ambroziak J, Henry SA. INO2 and INO4 gene products, positive regulators of phospholipid biosynthesis in Saccharomyces cerevisiae, form a complex that binds to the INO1 promoter. Journal of Biological Chemistry 1994;269:15344–9. 10.1016/S0021-9258(17)36612-7.

[35] Block-Alper L, Webster P, Zhou X, Supeková L, Wong WH, Schultz PG, et al. IN02, A Positive Regulator of Lipid Biosynthesis, Is Essential for the Formation of Inducible Membranes in Yeast. Mol Biol Cell 2002;13:40–51. 10.1091/mbc.01-07-0366.

[36] Carman GM, Han G-S. Regulation of phospholipid synthesis in yeast. J Lipid Res 2009;50:S69–73. 10.1194/jlr.R800043-JLR200.

[37] Du H-N, Fingerman IM, Briggs SD. Histone H3 K36 methylation is mediated by a *trans*-histone methylation pathway involving an interaction between Set2 and histone H4. Genes Dev 2008;22:2786–98. 10.1101/gad.1700008.

[38] Deshpande N, Bryk M. Diverse and dynamic forms of gene regulation by the S. cerevisiae histone methyltransferase Set1. Curr Genet 2023;69:91–114. 10.1007/s00294-023-01265-3.

[39] Ramakrishnan S, Pokhrel S, Palani S, Pflueger C, Parnell TJ, Cairns BR, et al. Counteracting H3K4 methylation modulators Set1 and Jhd2 co-regulate chromatin dynamics and gene transcription. Nat Commun 2016;7. 10.1038/ncomms11949.

[40] Krogan NJ, Kim M, Tong A, Golshani A, Cagney G, Canadien V, et al. Methylation of Histone H3 by Set2 in *Saccharomyces cerevisiae* Is Linked to Transcriptional Elongation by RNA Polymerase II. Mol Cell Biol 2003;23:4207–18. 10.1128/MCB.23.12.4207-4218.2003.

[41] Shilatifard A. The COMPASS Family of Histone H3K4 Methylases: Mechanisms of Regulation in Development and Disease Pathogenesis. Annu Rev Biochem 2012;81:65–95. 10.1146/annurev-biochem-051710-134100.

[42] Liu Z, Li X, Wang T, Zhang H, Li X, Xu J, et al. SAH and SAM/SAH ratio associate with acute kidney injury in critically ill patients: A case-control study. Clinica Chimica Acta 2024;553. 10.1016/j.cca.2023.117726.

[43] Basu Ball W, Neff JK, Gohil VM. The role of nonbilayer phospholipids in mitochondrial structure and function. FEBS Lett 2018. 10.1002/1873-3468.12887.

[44] Mejia EM, Hatch GM. Mitochondrial phospholipids: role in mitochondrial function. J Bioenerg Biomembr 2016. 10.1007/s10863-015-9601-4.

[45] Hermesh O, Genz C, Yofe I, Sinzel M, Rapaport D, Schuldiner M, et al. Yeast phospholipid biosynthesis is linked to mRNA localization. J Cell Sci 2014. 10.1242/jcs.149799.

[46] J.P. van Dijken JB, Brambillac L, Dubocd P, Francoise JM, Gancedof C, Giusepping, M.L.F.H, Eijnenh JJ, et al. An interlaboratory comparison of physiological and genetic properties of four Saccharomyces cerevisiae strains. Enzyme Microb Technol 2000;26:706–14.

[47] Rossmann MP, Stillman B. Immunoblotting Histones from Yeast Whole-Cell Protein Extracts. Cold Spring Harb Protoc 2013;8. 10.1101/pdb.prot067116.

[48] Yang S, Xue J, Ye C. Protocol for rapid and accurate quantification of phospholipids in yeast and mammalian systems using LC-MS. STAR Protoc 2022;3. 10.1016/j.xpro.2022.101769.

