## Supplementary Tables and Figures for "Gcn4 regulates phospholipid methylation and SAM dependent methyl allocations between phospholipids and histones during methionine-supplemented growth"

**Supplementary Table S1:** Strain list

| Strains | Genotype | Reference |
|---|---|---|
| WT (CEN.PK) | Mat a | This study |
| Gcn4-HA | MAT a, *GCN4-HA-KanMX* | This study |
| Ino2-HA | | This study |
| *gcn4Δ::NAT* | MAT a, *gcn4Δ::NAT* | This study |
| *gcn4Δ::KanMX* | MAT a, *gcn4Δ::KanMX* | This study |
| Opi3-HA | Mat a, *OPI3-HA-KanMX* | This study |
| Cho2-HA | Mat a, *CHO2-HA-KanMX* | This study |
| Psd1-HA | Mat a, *PSD1-HA-KanMX* | This study |
| *gcn4Δ::NAT* Cho2-HA | Mat a, *CHO2-HA-KanMX* | This study |
| *gcn4Δ::NAT* Opi3-HA | Mat a, *OPI3-HA-KanMX* | This study |
| *Ino2Δ::KanMX* Opi3-HA | Mat a, *OPI3-HA-KanMX* | This study |
| *gcn4Δ::NATIno2Δ::KanMX* Opi3-HA | Mat a, *OPI3-HA-Hyg* | This study |
| *Ino2Δ::KanMX* Cho2-HA | Mat a, *CHO2-HA-Hyg* | This study |
| *gcn4Δ::NATIno2Δ::KanMX* Cho2-HA | Mat a, *CHO2-HA-Hyg* | This study |

**Supplementary Table S2:** Primers used in this study

| Primer | Sequence |
|---|---|
| INO2 KO F1 | GTGCAATAAATAAATACATGGAACAGCAAAGGAGAAAATGcggatccccgggttaattaa |
| INO2 R1 | CGGGAGGCCATTTTCATCACTAATAGCTTGTATGAGCTCAgaattcgagctcgtttaaac |
| INO2 KO CHK Fw | ACACACCAGGACTCAAACAG |
| INO2 C' tag F2 | AAGCGCAAATGAAGCACTACAGCACATACTGGATGATTCCcggatccccgggttaattaa |
| INO2 C' tag CHK Fw | GGAGTTACCCACAAACACAG |
| CHO2 C' tag CHK Fw | AACAAGAGTTGGATCAGGTG |
| CHO2 C' tag F2 | TCGCGCTTGGGATATAAAACAAACGCTTGATAGTCTTGCTcggatccccgggttaattaa |
| CHO2 R1 | AGTACTTTTTAAATATATATACTCAAAAAAAAAAAACTCAgaattcgagctcgtttaaac |
| OPI3 C' tag CHK Fw | AACAGCCTACGTGTTCTCTG |
| OPI3 C' tag F2 | CATGATCTACGCTAACCGTGATAAGGCCAAAAAGAATATGcggatccccgggttaattaa |
| OPI3 C' tag R1 | GGCTTCTAACATTATAGAATATATAGAAATAGAGCACTTAgaattcgagctcgtttaaac |
| PSD1 C' tag CHK Fw | TGACTGGGTTTGTAAGGTTC |
| PSD1 C' tag F2 | GGGACAGAAATTAGGCATAATTGGAAAGAATGATTTAAAAcggatccccgggttaattaa |
| PSD1 R1 | ACAGCAAAATAAATGCTAACTTTACATATGATTGCTTTCAgaattcgagctcgtttaaac |

**Supplementary Table S3:** List of Q1/Q3 for metabolite (SAM and SAH) detection by LC-MS/MS.

| Sl no. | Metabolite | Q1/Q3 | CE | RT |
|---|---|---|---|---|
| 1 | S-Adenosyl Methionine (SAM) | 399/250 | 13 | 4.6 |
| 2 | S-Adenosyl Homoysteine (SAH) | 385/136 | 19 | 7.5 |

**Supplementary Table S3:** List of Q1/Q3 for metabolite (Phospholipids) detection by LC-MS/MS

| Sl no. | Metabolite | Q1 | Q3 | CE | RT |
|---|---|---|---|---|---|
| 1 | PC (16:1-16:1) | 788.545 | 253.217 | -50 | 16.2 |
| 2 | PC (16:1-18:1) | 816.576 | 281.249 | -50 | 20.4 |
| 3 | PE (16:1/16:1) | 686.476 | 253.217 | -50 | 9.7 |
| 4 | PE (16:1/18:1) | 714.508 | 281.249 | -50 | 10.4 |
| 5 | PS (16:1/16:1) | 730.476 | 253.217 | -50 | 7.2 |
| 6 | PS (16:1/18:1) | 758.498 | 253.217 | -50 | 7 |

Supplementary Figure 1

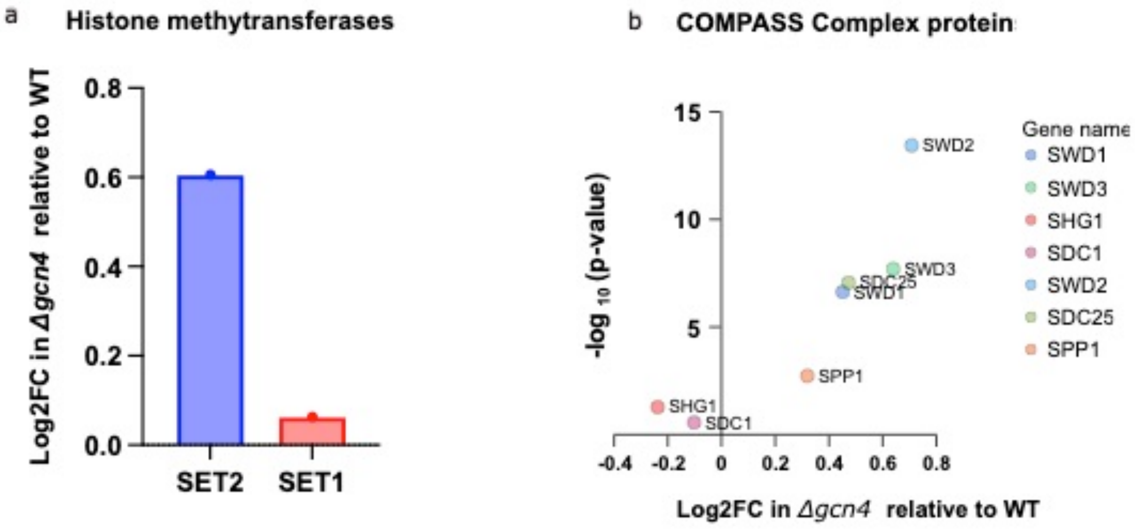


***Supplementary Figure 1: Differential expression of histone methyltransferases in the △gcn4 mutants relative to the WT cells grown in the medium supplemented with methionine*** *a.* Boxplot showing the log2 FC values of the Histone methyltransferases SET2 and SET1 in △*gcn4* relative to WT cells. This data is from methionine supplemented conditions. b. Volcano plot showing the relative fold change (Log2 scale) in the expression of COMPASS complex proteins in △*gcn4* deletion mutant relative to the wild type cells. Y-axis shows the significance of the differential expression as -log10(*p-value).*
